# Mapping Molecular Diversity in Prostate Cancer with a Combined Multiplex IHC and RNA-ISH Assay

**DOI:** 10.1101/2025.08.27.672653

**Authors:** Shannon Carskadon, Sean Williamson, Sangeetha Jyothilingam, Nilesh Gupta, Nallasivam Palanisamy

## Abstract

Combining multiplex Immunohistochemistry (mIHC) and RNA in situ hybridization (RNA-ISH) in a single assay allows for the simultaneous detection of multiple markers in tissue sections. This approach can be a powerful tool for understanding the molecular landscape of tumors, such as prostate cancer, by providing information in the same tissue context. Studying the expression of multiple genes in the same tissue is often necessary. ERG, SPINK1, ETV1, and ETV4 are all relevant genes in the context of prostate cancer. ERG is a transcription factor, and its fusion with the TMPRSS2 gene is one of the most common genetic alterations in prostate cancer. Overexpression of the ERG protein due to this gene, involved in gene fusion with TMPRSS2 and rarely with other 5’ partner genes, can be detected using Immunohistochemistry (IHC) due to the availability of a characterized antibody. SPINK1 (serine peptidase inhibitor, Kazal type 1) is an overexpressed protein in a subset of prostate cancers that do not harbor ERG rearrangements. High SPINK1 expression is known to be associated with a more aggressive tumor phenotype in some studies. SPINK1 expression can be detected using a well-characterized antibody by IHC. ETV1 and ETV4 are transcription factors that can also be involved in gene fusions in prostate cancer, albeit less frequently than ERG. There are no well-characterized antibodies for ETV1 and ETV4. Due to the mutually exclusive expression of ERG, SPINK1, ETV1, and ETV4, simultaneous evaluation of these genes can provide valuable information on the molecular subtype of prostate cancer, which can have implications for the assessment of inter- and intra-tumor heterogeneity, prognosis, and potential treatment. We describe a novel approach for the simultaneous evaluation of all four markers in prostate cancer tissues using a combined dual IHC and dual RNA-ISH method using validated antibodies for ERG, SPINK1, and RNAscope probes for ETV1 and ETV4.

## INTRODUCTION

Prostate cancer exhibit complex molecular and morphological heterogeneity. Presence of multiple ETS family related gene fusions and SPINK1 expression in a mutually exclusive pattern in a subset of prostate cancer offers technical challenges for reliable evaluation of these markers. Combining Dual Immunohistochemistry (Dual-IHC) with Dual RNA in situ hybridization (Dual-RNA-ISH) for the simultaneous evaluation of biomarkers in prostate cancer offers several significant advantages. The crucial advantage of this approach includes a holistic view of tumor biology by capturing the gene expression pattern in a single tissue sample. This combined approach provides a deeper understanding of the molecular mechanisms driving cancer. One can determine if the expressed gene is in the same cell population or expressed with other markers, which is vital in understanding the functional significance of multiple-driver molecular aberrations, particularly in prostate cancer, which exhibits a rare phenomenon of harboring multiple-driver molecular biomarkers within the multifocal disease. By examining ERG, SPINK1, ETV1, and ETV4 simultaneously, researchers and clinicians can classify tumors more accurately into distinct molecular subtypes, guiding tailored treatment strategies. Analyzing multiple markers in a single tissue section maximizes the use of often-limited availability tissue samples, especially when dealing with biopsy specimens. Combining Immunohistochemistry (IHC) and RNA in situ hybridization (RNA ISH) on the same tissue section is a valuable technique, especially when tissue samples are limited, as is often the case with biopsy specimens. RNA-ISH is particularly useful when no specific antibody is available for a gene of interes.[1-6].

Prostate biopsies and other tissue samples can be small, and conserving tissue is essential. Combining IHC with RNA-ISH allows for the simultaneous evaluation of multiple markers in a single tissue section, providing comprehensive molecular information without additional sections[1, 3]. Using the same tissue section for multiple marker screening reduces the number of slides and reagents needed as well as the time required for processing, leading to more efficient use of resources. Evaluating various markers on the same tissue section ensures that any observed molecular features or patterns genuinely represent that specific tissue area, reducing the risk of discrepancies from analyzing different tissue sections. For clinicians, having a comprehensive molecular profile from a single tissue section can aid in more informed decision-making regarding treatment strategies. A combined assay can provide a clearer picture of the expression pattern of multiple genes to understand their role in tumor development. Specific tumor subtypes or conditions may be rare in research settings, leading to scarce tissue samples. Making the most out of each tissue sample is imperative in such scenarios, and a combined approach will eliminate the discrepancies associated with tissue sections cut at different levels of the tissue blocks. Given the mutually exclusive expression patterns of ERG, SPINK1, ETV1, and ETV4, there are no technical challenges to optimizing the assay. Furthermore, mutually exclusive expression makes it easy to distinguish between them without any overlap or interference[7].

Having the ability to test multiple markers from the same sample can validate findings more robustly for researchers. ERG, SPINK1, ETV1, and ETV4 — and their cumulative contribution to about 60% of prostate cancers shed light on the molecular heterogeneity that characterizes prostate cancer. Their mutual exclusivity in expression underlines the complexity and diversity of tumor subtypes in this disease. The fact that these markers are expressed mutually exclusively suggests distinct molecular subclasses of prostate cancer. Different genetic alterations may drive each subtype, which pave the way for a more refined molecular classification of prostate cancer based on these markers. Each of these markers might correlate with distinct clinical outcomes. For instance, while ERG fusions have been extensively studied and have known associations with disease progression, other markers like SPINK1 overexpression might have unique prognostic implications. Understanding these distinctions can help in better risk stratification of patients. Identifying and validating these mutually exclusive markers suggests potential therapeutic targets. For example, tumors expressing a particular marker might respond more to specific targeted therapies, leading to the development of a more personalized treatment approach for prostate cancer patients. As our understanding deepens, the potential for developing tests that can quickly identify the molecular subtype of prostate cancer based on these markers becomes realistic. The molecular diversity of prostate cancer might also explain why some patients resist specific treatments. Some treatments might be effective against one subtype but not others. The >60% coverage by these markers still leaves a considerable proportion of prostate cancers uncharacterized at the molecular level. Further research into the remaining 40% might uncover additional significant markers or genetic alterations. While identifying these markers provides a solid foundation, translating these findings into clinical practice requires rigorous validation. It is essential to ensure that the detection methods are accurate and reproducible and that the clinical implications of each subtype in terms of prognosis, and treatment response are thoroughly understood. Identifying these mutually exclusive markers in prostate cancer has provided valuable insights into the molecular landscape of the disease. It emphasizes the need for a more personalized approach to diagnosis, prognosis, and treatment. The challenge lies in translating these findings into tangible patient benefits, which requires collaboration between researchers, clinicians, and the broader medical community.

## MATERIALS AND METHODS

### Study design and sample/patient selection

Tissue microarrays (TMAs) were generated from a cohort of 794 prostate cancer patients who underwent robotic radical prostatectomy at our institution, including 433 Caucasian (CA), 317 African American (AA), and 44 patients of other racial/ethnic backgrounds. For each patient, three tumor cores were sampled from distinct regions of the prostatectomy tissue blocks to account for intratumoral heterogeneity. Multiple TMA blocks were constructed, with each block representing tissue from 30 or more patients. Sections were cut from each TMA block and processed for downstream analyses as described below. The study protocol was approved by the Institutional Review Board (IRB).

### RNA in situ hybridization and triple Immunohistochemistry

We have developed methods for simultaneously evaluating the selected four markers by dual IHC and dual RNAISH methods [1]. IHC was used to detect ERG and SPINK1 simultaneously, and RNA-ISH (RNAscope) was selected for ETV1 and ETV4 due to the unavailability of cancer-specific antibodies[8]. For RNA-ISH, the slides were initially incubated at 60°C for 1 hour, then deparaffinized using xylene twice for 5 minutes each with periodic agitation. The slides were then immersed in 100% ethanol twice for 3 minutes, with occasional agitation, and air-dried for 5 minutes. After encircling the tissues with a pap pen and treating them with H_2_O_2_ for 10 minutes, the slides were rinsed in distilled water and boiled in 1X Target Retrieval for 15 minutes.

Subsequently, they were treated with Protease Plus for 15 minutes at 40°C in a HybEZ Oven (Bio-Techne, 310010). These steps were facilitated by the RNAscope Pretreatment kit (Bio-Techne, 310020). Following further rinses in distilled water, the slides were exposed to ETV1 (Bio-Techne, 311411) and ETV4 (Bio-Techne, 478571-C2) probes at a 50:1 ratio for 2 hours at 40°C in the HybEZ Oven. Two washes in 1X Wash Buffer (Bio-Techne, 310091) for 2 minutes each were performed, and the slides were stored overnight in a 5X SSC solution. The slides were washed twice in 1X Wash Buffer for 2 minutes each the next day. Then, they underwent treatment with Amp 1, Amp 2, Amp 3, and Amp 4 for specific durations at 40□C in the HybEZ oven, with two washes in 1X Wash Buffer for 2 minutes each after each step. Subsequently, the slides were treated with Amp 5 and Amp 6 for specified durations at room temperature in a humidity chamber, again with two washes in 1X Wash Buffer for 2 minutes each after each step. Red color development (ETV4) was achieved by adding a 1:60 solution of Fast Red B: Fast Red A to each slide and incubating for 10 minutes. The slides were washed twice in 1X Wash Buffer for 2 minutes each, then treated with Amp 7 and Amp 8 at specified durations at 40□C in the HybEZ oven, with two washes in 1X Wash Buffer for 2 minutes each after each step. Further treatment with Amp 9 and Amp 10 was performed for specified durations at room temperature in a humidity chamber, followed by two washes in 1X Wash Buffer for 2 minutes each after each step. Brown color development (ETV1) was achieved by adding a solution of Betazoid DAB (1 drop DAB to 1ml Buffer: Biocare Medical, BDB2004L) to each slide and incubating for 10 minutes. Amps 1-10 and Fast Red were (ETV4) included in the RNAscope 2.5 HD Duplex Detection Reagents (Bio-Techne, 322500). For IHC, slides were then washed in 1X EnVision FLEX Wash Buffer (DAKO, K8007) for 5 minutes. Subsequently, the slides were treated with Peroxidazed 1 (Biocare Medical, PX968M) for 5 minutes and Background Punisher (Biocare Medical, BP974L) for 10 minutes, with a wash of 1X EnVision FLEX Wash Buffer for 5 minutes after each step. Anti-ERG (EPR3864) rabbit monoclonal primary antibody (1:50; Abcam, ab92513) and a mouse monoclonal against SPINK1 (1:100; Novus Biologicals, H00006690-M01) were added to each slide, which was then cover-slipped with parafilm, placed in a humidifying chamber, and incubated overnight at 4□C. The following day, the slides were washed in 1X EnVision Wash Buffer for 5 minutes and then set in Mach2 Doublestain 1 (Biocare Medical, MRCT523L) for 30 minutes at room temperature in a humidifying chamber. Ferangi Blue solution (1 drop to 2.5ml buffer; Biocare Medical, FB813) was added and slides were incubated for 5 minutes. Then slides were washed in 1X EnVision FLEX Wash Buffer for 5 minutes and treated with a Vina Green solution (1 drop to 1ml buffer; Biocare Medical, BRR807) for 5 minutes. The slides were then rinsed twice in distilled water, followed by treatment with EnVision FLEX Hematoxylin (DAKO, K8008) for 2 minutes. After several rinses in tap water, the slides were dried completely. Finally, the slides were dipped in xylene approximately 15 times, and EcoMount (Biocare Medical, EM897L) was added to each slide, which was then cover-slipped. This protocol is not limited to the analysis of TMA alone, biopsy and wholemount sections of prostate can also be evaluated. It is important to note that RNA ISH should be performed first followed by IHC but not the otherway.

## RESULTS

A total of 794 prostate cancer cases were analyzed for the expression of ETS family gene fusions (ERG, ETV1, ETV4) and SPINK1. Of these, 386 cases (49%) demonstrated expression of a single marker, 28 cases (3.50%) expressed two different markers, and 380 cases (48%) were negative for all tested markers (Table 1).

**Table 1:** Distribution of molecular subtypes of prostate cancer by race based on ERG, SPINK1, ETV1, and ETV4 expression. In this cohort (n = 794), 49% of cases expressed at least one molecular marker, 3.5% showed dual marker expression, and 48% were negative for all tested markers. ERG expression was more common in Caucasian Americans compared to African Americans (29% vs. 20%, p = 0.004), whereas SPINK1 expression was more frequent in African Americans (23% vs. 13%, p < 0.001). Expression of ETV1 (4%) and ETV4 (2%) was infrequent across all groups and not significantly different by race. Co-expression of two markers was rare, with ERG+SPINK1+ being the most common combination (2%).

| Marker Status | Marker Expression | Caucasian American<br>(n=433) | African American<br>(n=317) | Other (n=44) | Total |
| --- | --- | --- | --- | --- | --- |
| Expression of a single marker (n=386, 48%) | ERG+ | 127 (29%)* | 63 (20%)* | 14 (32%) | 204 (26%) |
|  | SPINK1+ | 57 (13%)<sup>1</sup> | 74 (23%)<sup>1</sup> | 5 (11%) | 136 (17%) |
|  | ETV1+ | 20 (5%)<sup>2</sup> | 7 (2%)<sup>2</sup> | 5 (11%) | 32 (4%) |
|  | ETV4+ | 9 (2%)<sup>3</sup> | 4 (1%)<sup>3</sup> | 1 (2%) | 14 (2%) |
| Expression of two different markers (n=28, 3.5%) | ERG+ SPINK+ | 5 (1%) | 10 (3%) |  | 15 (2%) |
|  | ERG+ ETV1+ | 4 (1%) |  | 1 (2%) | 5 (0.6%) |
|  | ERG+ ETV4+ | 2 (0.5%) |  |  | 2 (0.3%) |
|  | SPINK+ ETV1+ | 3 (0.7%) | 1 (0.3%) |  | 4 (0.5%) |
|  | SPINK+ ETV4+ |  | 2 (0.6%) |  | 2 (0.3%) |
| Negative for all tested markers (n=380, 48%) |  | 206 (48%)<sup>□</sup> | 156 (50%)<sup>□</sup> | 18 (41%) | 380 (48%) |
|  | * p=0.004 | <sup>1</sup> p<0.001 | <sup>2</sup> p=0.12 | <sup>3</sup> p=0.57 | <sup>□</sup> p=0.71 |

### Expression of a Single Marker

Among tumors with a single positive marker, ERG was the most frequently detected alteration (n = 204, 26%), followed by SPINK1 (n = 136, 17%), ETV1 (n = 32, 4%), and ETV4 (n = 14, 2%). ERG expression was significantly more frequent in Caucasian Americans (127/433; 29%) compared with African Americans (63/317; 20%, p = 0.004). Conversely, SPINK1 expression was more common in African Americans (74/317; 23%) than in Caucasians (57/433; 13%, p < 0.001). ETV1 and ETV4 expression were rare and did not differ significantly between the two groups (p = 0.12 and p = 0.57, respectively).

### Co-expression of Two Markers

Dual expression of two distinct markers was observed in 28 cases (3.5%). The most frequent combination was ERG+SPINK1+ (n = 15), which was enriched in African American patients (10/317; 3%) compared with Caucasian Americans (5/433; 1%). Other co-expression patterns included ERG+ETV1+ (n = 5), ERG+ETV4+ (n = 2), SPINK1+ETV1+ (n = 4), and SPINK1+ETV4+ (n = 2). These dual-positive cases were rare across all racial groups.

### Negative for All Markers

Nearly half of the cohort (n = 380, 48%) was negative for ERG, SPINK1, ETV1, and ETV4. The distribution of marker-negative tumors was comparable between Caucasian Americans (48%) and African Americans (50%), with no significant difference (p = 0.71).

Taken together, these findings highlight significant racial differences in molecular subtypes of prostate cancer: ERG expression is enriched in Caucasian Americans, whereas SPINK1 expression is more prevalent in African Americans. Dual marker expression was uncommon but occurred more often in African Americans, particularly the ERG+SPINK1+ subtype.

### Biomarker Expression Across Pathological Categories

We next evaluated biomarker expression in relation to cancer status and Gleason grade groups (Table 2). Among the 38 benign cores, none expressed ERG, SPINK1, ETV1, or ETV4, whereas precursor lesions (HGPIN, n = 26) showed infrequent expression of ERG (n = 3), SPINK1 (n = 2), ETV1 (n = 2), and ETV4 (n = 1). Atypical cores (n = 2) rarely expressed markers (one case positive for SPINK1).

**Table 2:** Expression of molecular markers by cancer status and Gleason grade group. A total of 794 prostate tissue cores were evaluated. ERG expression was most frequent (25.8%), followed by SPINK1 (17.2%), ETV1 (3.9%), and ETV4 (1.6%). Dual expression of markers occurred in 3.6% of cores, most commonly ERG+SPINK1+ (1.9%). Nearly half of all cores (47.8%) were negative for all tested markers. ERG and SPINK1 positivity was observed across all Gleason grade groups, with ERG expression most frequent in Gleason Grade Group 2 (30% of all ERG+ cases) and SPINK1 enriched in Grade Groups 2–4. ETV1 and ETV4 were rare events, distributed across intermediate and high-grade cancers. Benign and HGPIN samples occasionally harbored marker expression, though less frequently than cancer samples.

| Cancer Status | ERG+ | SPINK1+ | ETV1+ | ETV4+ | ERG+ SPINK+ | ERG+ ETV1+ | ERG+ ETV4+ | SPINK+ ETV1+ | SPINK+ ETV4+ | Negative | Total Number |
| --- | --- | --- | --- | --- | --- | --- | --- | --- | --- | --- | --- |
| Benign |  |  |  |  |  |  |  |  |  | 38 | 38 |
| HGPIN | 3 | 2 | 2 | 1 |  |  |  |  |  | 18 | 26 |
| Atypical |  | 1 |  |  |  |  |  |  |  | 1 | 2 |
| Gleason Grade Group 1 | 55 | 17 | 3 | 1 | 1 | 2 |  |  |  | 66 | 145 |
| Gleason Grade Group 2 | 89 | 60 | 15 | 3 | 11 | 2 | 1 | 3 | 1 | 114 | 299 |
| Gleason Grade Group 3 | 29 | 21 | 6 | 5 | 3 | 1 | 1 |  |  | 60 | 126 |
| Gleason Grade Group 4 | 18 | 25 | 1 | 4 |  |  |  | 1 | 1 | 54 | 104 |
| Gleason Grade Group 5 | 10 | 10 | 5 |  |  |  |  |  |  | 29 | 54 |
| Total number of cores | 204 | 136 | 32 | 14 | 15 | 5 | 2 | 4 | 2 | 380 | 794 |
| Percentage | 25.8% | 17.2% | 3.9% | 1.6% | 1.9% | 0.6% | 0.3% | 0.5% | 0.3% | 47.8% | 100.0% |

### Gleason Grade Group 1–2

In low-grade cancers, ERG was the most frequently observed alteration. ERG expression was detected in 55/145 (38%) of Gleason grade group 1 tumors and 89/299 (30%) of Gleason grade group 2 tumors. SPINK1 expression was observed in 17/145 (12%) and 60/299 (20%) of grade groups 1 and 2, respectively. ETV1 and ETV4 expression were less common, but dual expression of ERG+SPINK1+ occurred in 1% of grade group 1 and 3.7% of grade group 2 cases (Table 2).

### Gleason Grade Group 3–5

In higher-grade tumors, expression of ERG decreased in frequency, being present in 29/126 (23%) of Gleason grade group 3, 18/104 (17%) of Gleason grade group 4, and 10/54 (19%) of Gleason grade group 5 cases. In contrast, SPINK1 expression was relatively stable or slightly enriched in higher grades, occurring in 21/126 (17%), 25/104 (24%), and 10/54 (19%) of grade groups 3, 4, and 5, respectively. ETV4 expression was more frequent in grade groups 3 and 4 compared with lower grades (5/126 (3.9%) and 4/104 (3.8), respectively. There was no difference in ETV1 (6/126) (4.7%) expression compared to ETV4.(Table 2).

### Negative Cases

Overall, 380/794 (48%) of tumors were negative for all tested markers. The proportion of marker-negative tumors increased with grade, from 46% in grade group 1 to 54% in grade groups 4 and 5.

These findings indicate that ERG expression predominates in lower-grade tumors and decreases in higher grades, whereas SPINK1 expression is more evenly distributed across grades and may be relatively enriched in Gleason grade groups 4–5. ETV1 and ETV4 alterations remain rare across all groups but appear slightly more common in intermediate- to high-grade cancers.

Overall, the results indicate that evaluation of these markers using our new approach on wholemount radical prostatectomy tissues and saturation biopsy samples will reveal even more molecular heterogeneity and identify new distinct molecular subtypes of prostate cancer.

## DISCUSSION

Prostate cancer, like many other malignancies, exhibits extensive molecular heterogeneity[4]. The individual or mutually exclusive expression of markers such as ERG, SPINK1, ETV1, and ETV4 in prostate tumors underscores this diversity[2, 3]. As our understanding of prostate cancer deepens, it is clear that molecular classification is as crucial as histological grading. Molecular subtypes might provide insights into the etiology and progression mechanisms and offer refined prognostic and therapeutic avenues. Traditional pathology heavily relies on histology and IHC for diagnosing and classifying tumors. However, the advent of molecular techniques, including RNA ISH, has brought forward an additional layer of data that could be pivotal for a holistic understanding of tumor biology. By leveraging both proteins and RNA from the same tissue, there is an opportunity to bridge the gap between genotypic variations and phenotypic manifestations in prostate cancer. The direct comparison and correlation between gene expression and protein presence in the same cellular environment can shed light on post-transcriptional and translational events, potentially highlighting therapeutic targets or resistance mechanisms[5].

Biopsy samples, especially from deep-seated tumors or patients undergoing multiple biopsies, can be limiting. Each tissue section becomes a valuable information repository; thus maximizing data retrieval from every section becomes crucial. The combined dual IHC and dual RNA ISH approach ensures that researchers and clinicians glean maximal molecular data without additional tissue sections, preserving samples for other potential investigations. This combined method offers a more exhaustive molecular profiling approach in research, especially when investigating rare tumor subtypes or when tissue samples are hard to procure. Moreover, from a clinical standpoint, a comprehensive molecular snapshot can significantly influence therapeutic decisions, especially in the era of precision medicine. While the combined approach is promising, it is challenging. The future will see more integrated diagnostic approaches, where multiple molecular layers are evaluated concurrently, paving the way for genuinely personalized therapeutic strategies. In conclusion, prostate cancer’s molecular complexity necessitates innovative approaches to understanding and management. Integrating IHC and RNA ISH is a significant step in this direction, offering a nuanced view of tumor biology while conserving invaluable tissue samples.

In this study, we comprehensively analyzed the expression of ETS family gene fusions (ERG, ETV1, ETV4) and SPINK1 across a racially diverse cohort of prostate cancer patients. Our findings underscore the molecular heterogeneity of prostate cancer and demonstrate both racially-associated molecular subtypes and grade-associated expression patterns.

Consistent with prior reports, ERG positivity was significantly enriched in Caucasian American patients, whereas SPINK1 expression was more prevalent in African American patients. These findings reinforce the concept that prostate cancer subtypes are not uniformly distributed across racial groups and may contribute to observed disparities in clinical presentation and outcomes. Importantly, while dual-marker expression was rare overall, the ERG+SPINK1+ subtype occurred more frequently in African Americans, suggesting that molecular overlap between canonical drivers may be more common in this group. These observations highlight the importance of considering racial background in molecular profiling and biomarker-driven patient stratification.

When analyzed in the context of tumor grade, distinct patterns emerged. ERG expression was highest in lower-grade tumors (Gleason grade groups 1–2) and declined significantly with increasing Gleason grade, as confirmed by both logistic regression and Cochran–Armitage trend analyses. This pattern suggests that ERG-driven tumors may represent a biologically distinct subset enriched in early, lower-grade disease. In contrast, SPINK1 expression was relatively stable across grade groups, with a non-significant trend toward enrichment in higher-grade tumors (particularly grade groups 4–5). This observation aligns with prior studies associating SPINK1 with aggressive clinical features, although our findings did not reach statistical significance. Expression of ETV1 and ETV4 remained infrequent across all grade groups, but appeared slightly more common in intermediate- to high-grade cancers, consistent with their role as less prevalent but biologically relevant drivers.

Approximately half of all tumors in this cohort were negative for ERG, SPINK1, ETV1, and ETV4, highlighting the existence of additional, hitherto unidentified molecular alterations that drive prostate cancer. This large negative subgroup suggests that molecular classification based solely on these canonical markers is incomplete, and emphasizes the need for expanded genomic profiling approaches using next-generation sequencing and spatial transcriptomics to capture the full spectrum of disease heterogeneity.

Taken together, these findings have important implications. First, they provide further evidence that molecular subtypes of prostate cancer differ by race, a factor that should be integrated into biomarker-driven clinical trial design and precision oncology approaches. Second, they show that ERG- and SPINK1-defined tumors follow divergent trajectories across Gleason grades, supporting their use not only as molecular classifiers but also as potential prognostic indicators. Finally, the large fraction of marker-negative tumors highlights a critical gap in our current understanding of prostate cancer biology, warranting further investigation into alternative drivers and pathways.Further investigations, using wholemount radical prostatectomy and satutation biopsy samples using this novel approach will reveal futher the genetic complexity of prostate cancer.

## Acknowledgement

We thank Natalia Draga, Jingli Yang for their help in the preparation of slides from biopsy blocks. This study was supported by a US Department of Defense grant W81XWH-16-1-0544 to Nallasivam Palanisamy.

## Figures

**Figure 1:**
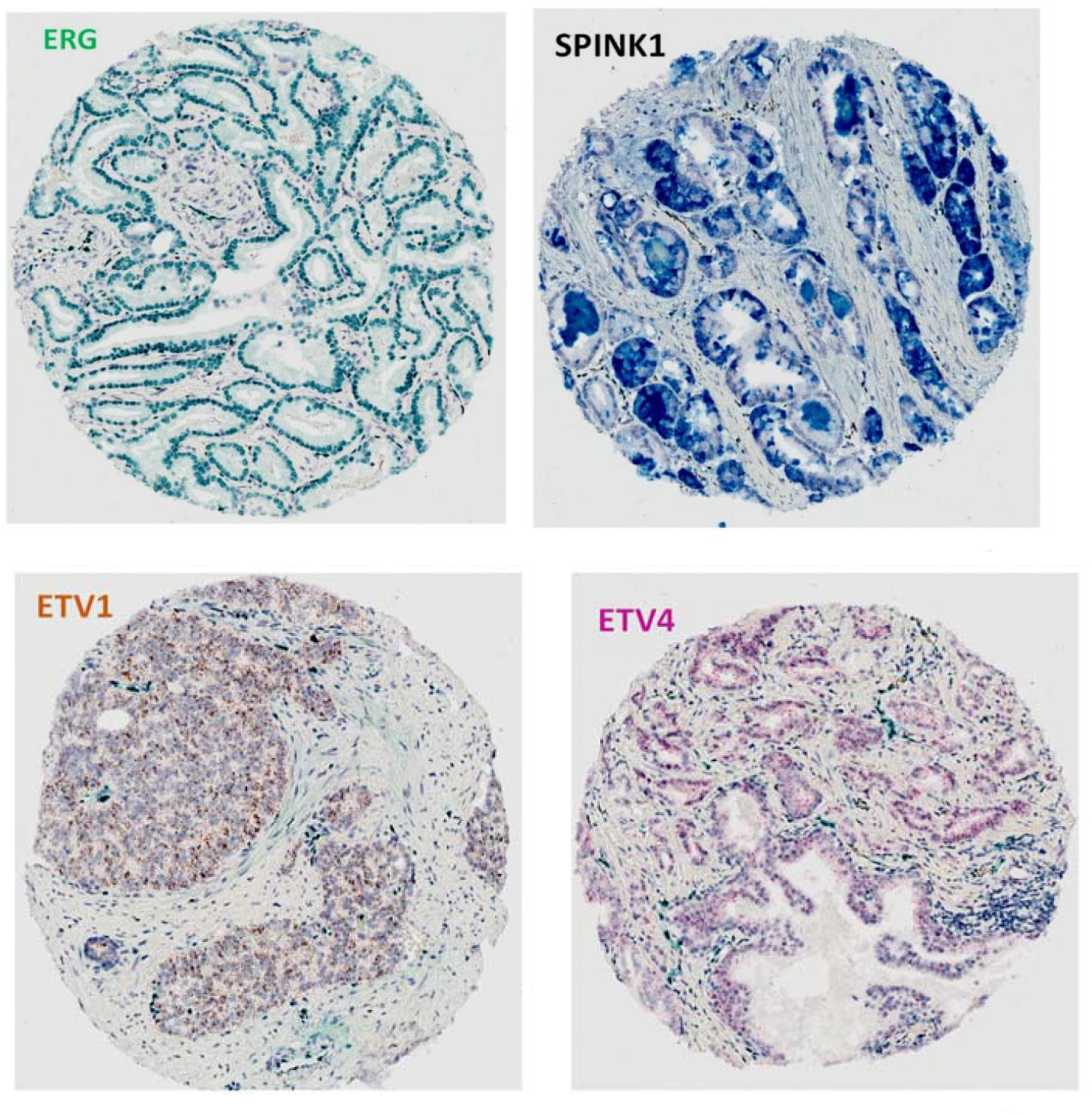
Expression of ERG, SPINK1, ETV1, and ETV4 in prostate tissue. Representative tissue microarray cores showing molecular marker expression in prostate samples. ERG expression (green, top left) highlights nuclear staining in epithelial cells. SPINK1 expression (blue, top right) demonstrates cytoplasmic staining in a distinct subset of tumor cells. ETV1 (brown, bottom left) and ETV4 (magenta, bottom right) show cytoplasmic staining patterns (punctate dots corresponding to RNA transcript), respectively, in discrete tumor regions. These images illustrate the heterogeneous expression of ETS family genes and SPINK1 in prostate cancer tissues.

**Figure 2:**
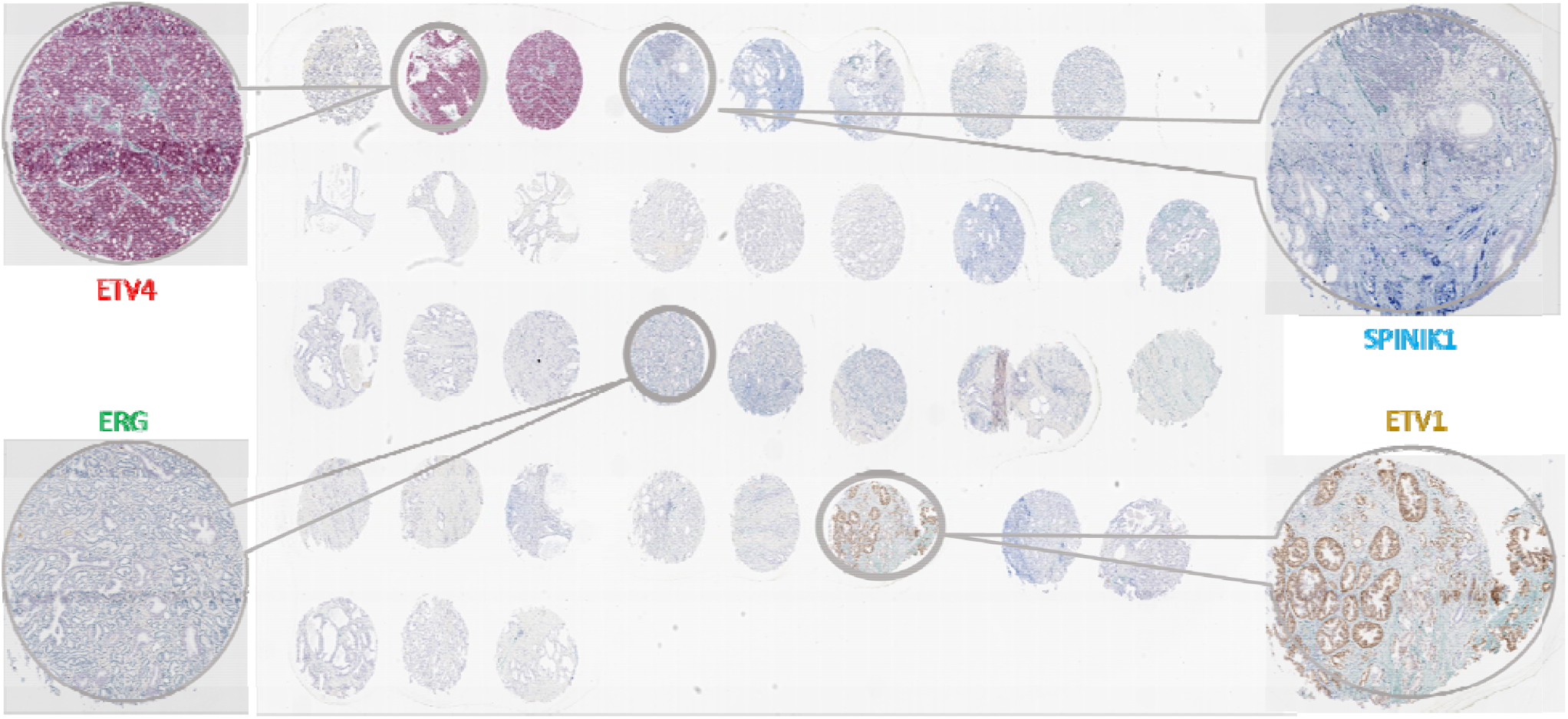
Representative multiplex analysis of molecular subtypes in prostate cancer tissue microarrays. Tissue microarray (TMA) cores were subjected to a combined assay utilizing dual immunohistochemistry (IHC) for ERG and SPINK1 and dual RNA in situ hybridization (RNA-ISH) for ETV1 and ETV4, enabling simultaneous detection of protein and RNA markers within the same tissue section. The central panel shows a TMA with multiple prostate cancer cores, with magnified insets highlighting distinct molecularly defined tumor subsets: ERG-positive (green), SPINK1-positive (blue), ETV1-positive (brown), and ETV4-positive (red) tumors. Each marker exhibits mutually exclusive expression patterns, underscoring the molecular heterogeneity of prostate cancer. Insets illustrate the specific staining patterns for each marker at higher magnification, demonstrating the successful application of combined IHC and RNA-ISH for multiplex molecular subtyping in prostate cancer.

